# *Nematocida displodere* Mechanosensitive Ion Channel of Small Conductance 2 assembles into a unique 6-channel super-structure *in vitro*

**DOI:** 10.1101/2024.03.27.587072

**Authors:** Alexandra Berg, Ronnie P-A Berntsson, Jonas Barandun

## Abstract

Mechanosensitive ion channels play an essential role in reacting to environmental signals and sustaining cell integrity by facilitating ion flux across membranes. For obligate intracellular pathogens like microsporidia, adapting to changes in host environment is crucial for survival and propagation. Despite representing a eukaryote of extreme genome reduction, microsporidia have expanded the gene family of mechanosensitive ion channels of small conductance (*mscS*) through repeated gene duplication and horizontal gene transfer. All microsporidian genomes that are characterized to-date contain *mscS* genes of both eukaryotic and bacterial origin, and have at least 5 different *mscS* copies. Here, we investigated the cryo-electron microscopy structure of the bacterially derived mechanosensitive ion channel of small conductance 2 (MscS2) from *Nematocida displodere*, an intracellular pathogen of *Caenorhabditis elegans*. MscS2 is the most compact MscS known, and assembles into a unique superstructure *in vitro* with six heptameric MscS2 channels oligomerizing through their transmembrane domains. Individual MscS2 channels are oriented in a heterogeneous manner to one another, resembling an asymmetric, flexible six-way cross joint. Finally, we show that, despite the extreme compaction, microsporidian MscS2 still forms a heptameric membrane channel, conserving the most important structural features of bacterial MscS.

## Introduction

Microsporidia compose a large group of highly adapted, obligate intracellular pathogens that modified their genomic repertoire for exceedingly efficient parasitism (1,2). Their early divergence within the fungal kingdom and their development at a fast evolutionary rate (3), shaped a highly divergent phylum of unicellular eukaryotes (4). The phylum of microsporidia infects a broad range of hosts, covering almost all animal taxa including humans (5–7). While generally ubiquitous, microsporidia thrive in agriculturally kept aquatic and terrestrial animals (6), where mass rearing, antibiotic and pesticidal overuse may benefit parasite transmission in immunocompromised and stressed hosts (8–10). Affected animals are *e.g.* shrimp, salmon, and honeybees, which are important contributors to the global food supply. Microsporidian infections can therefore lead to significant economic losses (8). In humans, microsporidia can cause disease in otherwise healthy individuals, but mainly cause opportunistic infections in immunocompromised hosts, such as AIDS patients, organ transplant recipients and cancer patients (6,11,12).

The onset of an infection is typically initiated through the ingestion of dormant microsporidian spores by a host from contaminated sources (8,12). The spores migrate to the gastrointestinal tract and germinate in response to an appropriate stimulus (13,14). Spore germination triggers the explosive shoot out of the unique invasion organelle, the polar tube, through which the infectious cell content, the sporoplasm, is rapidly transported and injected into a host cell (15,16). Intracellularly, the sporoplasm hijacks host nutrients (17–19) which is vital for differentiation into a multinucleated, proliferating cell, the meront. The meront further develops into a sporont, that undergoes cellularization and synthesizes the spore coat, followed by the sporoblast stage, in which organelle formation takes place. Finally, the sporoblast evolves into a mature spore which can start a new infection cycle (20).

The intricate and successful nature of microsporidian parasitism is a result of efficient genome organization and compaction which makes them highly dependent on their host for replication and development (21–23) but for example nearly halves their energy requirements in form of ATP (24). To achieve this, microsporidia minimized or removed many enzymes involved in biosynthetic or metabolic pathways whose functions can be compensated through host system exploitation (2,18,21–23). Proteins required to exploit the host, but also to escape its immune system, were gained through horizontal gene transfer (HGT) from bacteria (2), or the host (25,26) and modified via gene duplications (26).

One universally conserved protein family, that has undergone an unusual evolution in microsporidia, are the mechanosensitive ion channels of small-conductance (MscS). MscS are membrane-embedded homo-heptameric proteins that exist in both eukaryotes and procaryotes and are best studied in *Escherichia coli*. Their main function as “osmotic safety valves” prevents the cell from bursting during hypoosmotic shock (27). There, a reduction in extracellular osmolarity induces swelling of the bacterial cell, consequently stretching the membrane. MscS opens in response to membrane tension (28) and enables the efflux of water and selected ions to release turgor pressure and restore cellular homeostasis (27). Similarly, in *Saccharomyces pombe* the MscS-like channels Mys1 and Mys2 are involved in the hyperosmotic shock response (29).

Microsporidian genomes harbor at least five copies of *mscS* (2) which, according to phylogenetic analyses (2), can be distinguished into two subfamilies: MscS1, derived from eukaryotes through lineage-specific expansion, and MscS2, originating from bacteria and likely acquired via HGT by the last common ancestor of microsporidia (2). While *mscS1* is represented with ≥ 4 copies in each microsporidian genome sequenced to date, only a single copy of *mscS2* can be identified. Further, MscS1 resembles a canonical, eukaryotic MscS in size and domain architecture, whereas MscS2 is substantially truncated in primary sequence; a consequence of losing one of the two C-terminal domains and most transmembrane domains (TMDs) (2,30), relative to the bacterial MscS. Of the three TMDs in bacterial MscS, the third (TMD3) is essential for ion channel function (31). Thus, the observed compaction phenomenon of the microsporidian MscS2 makes this evolutionary outlier an interesting target to study structure, oligomerization, channel formation, or membrane interaction.

To understand the impact of compaction on the structure and function of microsporidian MscS2, we determined a low-resolution cryo-EM structure of MscS2 from *Nematocida displodere*, an intracellular pathogen of *Caenorhabditis elegans*. We show that MscS2 forms a homo-heptameric channel that maintains only the essential domains (31–33) required for mechanosensitive ion channel function. Under our *in vitro* conditions, the variant produces a higher order superstructure, with six heptamers together forming a structure that resembles an asymmetric, flexible six-way cross joint. While the observed superstructure likely is the result of applied experimental procedures and represents a physiologically non-relevant assembly, it serves as a unique and intriguing model that challenges our understanding of protein-complex formation. In addition, it shows that the minimal microsporidian MscS2 can still form a heptameric, membrane-bound channel with a central pore.

## Materials and methods

### Bioinformatics analyses

#### Multiple sequence alignment

Microsporidian MscS2 were identified through blasting against the non-redundant NCBI database, against one at the time unpublished microsporidian genome of *Vairimorpha necatrix* (34) and against a local database created from three available, but non-annotated genomes of *Antonospora locustae* (35–37). Two multiple sequence alignments of 35 identified sequences with and without *E. coli* MscS were created in ClustalW (accessed September 2023) (38) with default settings and colored using Clustal X default Coloring in Jalview (v2.11.2.6) (39).

#### Cladogram generation

35 MscS2 sequences were aligned using MUSCLE (v5.1) (38) and trimmed with trimAl (v1.4.1) (40) (to remove spurious sequences and poorly aligned regions from a multiple sequence alignment). The resulting sequences were used to generate a cladogram with IQ-TREE (v2.0.3) (41,42) with 1000 bootstrap replicates and the MFP option for choice of substitution model.

Sequences used for the multiple sequence alignment and cladogram: *Annicaliia algerae, Antonospora locustae, Dictyocoela muelleri, Edhazardia aedis, Encephalitozoon cuniculi, Encephalitozoon hellem, Encephalitozoon intestinalis, Encephalitozoon romaleae, Enterocytozoon bieneusi, Enterocytozoon hepatopenaei, Enteropsecta breve, Enterospora canceri, Hamiltosporidium magnivora, Hamiltosporidium tvaerminnensis, Hepatospora eriocheir, Nematocida ausubeli, Nematocida displodere, Nosema granulosis, Nematocida homosporus, Nematocida major, Nematocida parisii, Nematocida species ERTm2, Nematocida species ERTm5, Ordospora colligata, Ordospora pajunii, Pancytospora epiphaga, Pancytospora philotis, Spraguea lophii, Thelohania contejeani, Trachipleistophora hominis, Tubulinosema ratisbonensis, Vairimorpha ceranae, Vairimorpha ceranae BRL01, Vairimorpha necatrix, Vavraia culicis*.

#### Topology prediction

For transmembrane region and signal peptide predictions of the 35 MscS2, we used DeepTMHMM (v1.0.24) (43) and TOPCONS (v2.0) (44). All sequences were entered individually to obtain a result graph for each protein.

#### Hydropathicity prediction

To identify primary-sequence based hydrophobic regions in *N. displodere* MscS2, we employed ProtScale on the ExPASy server (45) to predict the hydropathicity (46) of the protein.

#### AlphaFold prediction

The AlphaFold models for *N. displodere MscS2* were generated with ColabFold (v1.5.5) (47). First, a single-chain model was predicted using the default model type alphafold2_ptm. The heptamer model was created using the first rank monomer as template, with alphafold2_multimer_v3 model type. For both predictions we employed 20 prediction cycles and an early stop tolerance of 0.2. To display heptamer model confidence, we colored the model according to the pLDDT (predicted local-distance difference test) confidence measure (48).

### Cloning, protein production and bacterial cell lysis

We ordered *E. coli* codon-optimized *N. displodere mscS2* from Genewiz (Azenta Life Sciences) in a pUC-GW-Kan vector. Using primers listed in **S1 Table**, we amplified *N. displodere mscS2, mscS2Δ2-11* and *mscS2Δ2-29* via polymerase-chain reaction and cloned each gene into a *pRSFDuet* vector with an N-terminal His14-SUMO tag and a kanamycin resistance cassette. We transformed chemically competent *E. coli TOP10* cells with each plasmid, respectively, for propagation, validated positive clones using colony-PCR and sequencing. Subsequently, we transformed *E. coli* Rosetta (DE3) cells with the vector. For protein production, we inoculated 100 mL Luria Bertani (LB) medium containing 50 μg/mL kanamycin and 33 μg/mL chloramphenicol with colonies carrying the respective plasmid and grew them overnight at 37 °C with shaking. Next, we inoculated the main culture 1:35 in LB and further incubate culture at 37 °C and 150 rpm. When reaching OD600nm 0.7, the temperature was shifted 18 °C and induced expression by the addition of 1 mM IPTG for 18 h. The cells were then pelleted, and frozen in liquid nitrogen, before being lysed using a cryo-mill (Retsch), with three cycles of milling. The resulting bacterial powder was stored at −80 °C.

### Protein Purification

To purify MscS2Δ2-29, we thawed 5 g of the bacterial powder and resuspended it in 20 mL buffer A [50 mM HEPES-KOH, 300 mM NaCl, 10% glycerol, pH 8 at 4 °C], added 40 mM imidazole, 1 mM DTT, 1 mM MgCl_2_, DNase and protease inhibitors Phenylmethanesulfonyl Fluoride [10-20 µM], Pepstatin [1-2 µM] and E64 [1-2 µM]. We used a tissue grinder (Dounce homogenizer) to homogenize the suspension, before adding 1.5% (w/v) n-Dodecyl-beta-Maltoside (DDM) and incubating 1 h at 4 °C for solubilization. The insoluble fraction was removed by spinning down the sample at 48,500 x g for 1 h at 4 °C. The supernatant was loaded on a His-Trap 5 mL Ni-NTA column (Amersham Biosciences), washed three times with 10 column volumes buffer A containing 60 mM, 80 mM and 100 mM imidazole and 0.05% (w/v) DDM, respectively. The protein was eluted by buffer A containing 500 mM imidazole and 0.05% (w/v) DDM. Purification fractions were analyzed by sodium dodecyl sulfate polyacrylamide gel electrophoresis (SDS-PAGE). Next, to cleave the His14-SUMO tag we pooled all elution fractions that contained His14-SUMO-MscS2Δ2-29, transferred it to a 7 kilodalton cut-off SnakeSkin^TM^ dialysis tubing (Pierce^TM^, VWR), added Ulp1 protease [0.5 μM final concentration] and let it dialyze overnight at 4 °C in buffer A (with 7% glycerol) and with stirring. We assessed the successful cleavage via SDS-PAGE and split the sample in two. We analyzed one half using size-exclusion chromatography and dialyzed the other half again in buffer A without glycerol and with 0.05% DDM, to prepare MscS2Δ2-29 for cryo-EM grid freezing.^-^ For gel filtration experiments of MscS2Δ2-29 we used a Superdex 200 Increase 10/300 GL (Cytiva) set up on an ÄKTA^TM^ pure (Cytiva). We equilibrated the column with buffer A containing 0.05% DDM and 1 mM DTT without glycerol and loaded 500 μL pooled and cleaved elution sample.

### Negative-stain transmission electron microscopy

MscS2Δ2-29 was stained with 1.5% uranyl-acetate (EMS), as follows: We glow-discharged copper grids (200 squared mesh) coated with a thin carbon film for 30 sec at 15 mA, applied 3 μL of sample (concentration of 0.194 mg/mL), let it air dry for 30 sec, blotted off excess sample with Whatman paper, washed three times in 20 μL-drops of MiliQ water, stained for 30 sec in a 20μL-drop of uranyl acetate and remove excess stain with Whatman paper. Prior to storage at room temperature, we let the sample air dry for ∼15 minutes. We analyzed the MscS2Δ2-29 grids using a FEI Talos L 120C transmission electron microscope (Thermo Fisher Scientific), operating at 120 kV, and acquired TEM micrographs at a magnification of 73,000 with a Ceta 16M CCD camera employing the TEM Image & Analysis software (v4.17) (FEI).

### Cryo-EM grid preparation and data collection

For cryo-EM analysis of MscS2Δ2-29, we applied 4 μL aliquots (0.33 mg/mL) to glow-discharged (30 sec, 15 mA) Quantifoil R2/2 200-mesh copper grids with 2 nm carbon (EM sciences) at 4 °C and 100% humidity using the FEI Vitrobot Mark IV (Thermo Fisher Scientific). The sample was blotted for 5s, with −5 blot force prior to vitrification in liquid ethane.

Cryo-EM data were collected at the Umeå Core Facility for Electron Microscopy employing a 200 kV Glacios system (Thermo Fisher Scientific) equipped with a Falcon 4i direct electron detector. For MscS2Δ2-29, a total of 2000 movies, each with 40 frames, at a total dose of 50 e-/Å2, and a pixel size of 0.75 Å were collected using EPU (Thermo Fisher Scientific). We summarized the data collection statistics in S2 Table.

### Cryo-EM data processing

We processed the MscS2Δ2-29 cryo-EM data with Relion version 3.1 (47) and 4.0 (48). Movie alignment, drift correction and dose weighing were performed using MotionCor2 (49) and CTF estimation was done with CTFFIND (v4.1.14) (50). We manually inspected and removed micrographs with poor CTF or low ice quality resulting in 1856 micrographs from which 464,200 particles were extracted with a box size of 300 px using Laplacian autopicking in Relion. Particles were filtered using 2D classification followed by consensus refinement of 300,757 particles without imposed symmetry and resulting in a 6-channel superstructure of 8.27 Å resolution. A processing scheme is illustrated in **S1 Fig**.

### Mass photometry

For mass analysis, we used a RefeynMP2 instrument. We first cleaned a microscope cover slip (No. 1.5, 24×50 mm, Thorlabs) through alternated rinsing with MiliQ H_2_O and isopropanol, finishing off with MiliQ H_2_O and dried them under a clean nitrogen stream. On the center cover slip, we applied a silicone gasket, a self-adhesive frame with culture wells, to hold the later droplet sample. We diluted MscS2Δ2-29 samples in buffer [50 mM HEPES-KOH, 300 mM NaCl, pH 8 at 4 °C] to concentrations of 28 nM and 56 nM immediately before measurements. Buffer only was used as control. Each sample was applied to a fresh well, the focal position was determined, and movies of 60 sec duration were recorded. For data acquisition and evaluation, we used Refeyn Acquire MP and DiscoverMP software and NativeMarkTM (ThermoFisher) was used as standard.

## Results and Discussion

### Microsporidia retain the essential domains for ion-channel functionality in MscS2

We used different bioinformatics tools to identify MscS2 across microsporidia, obtain phylogenetic information, and predict domains and features to analyze the impact of compaction on structure and potential function. We identified 35 MscS2 through multiple rounds of blasting against the non-redundant NCBI database as well as against one at the time unpublished microsporidian genome from *Vairimorpha necatrix* (34) and against a local database created from three available, but non-annotated genomes of *Antonospora locustae* (35–37).

A multiple sequence alignment of these 35 MscS2 confirmed earlier findings that the MscS β-domain is conserved among microsporidia (**Fig 1A**). Residues located in important structural positions in the β-domain, e.g., the kink-inducing P129, G143, P166, are conserved between microsporidia and *E. coli* (**S2A Fig**) suggesting a similar tertiary fold. Further, the region just before the MscS β-domain, which in *E. coli* MscS (*Ec*MscS) corresponds to the amphipathic transmembrane helix (TMH) 3b, also shows reasonably high conservation (**Fig 1A and S2A Fig**). To identify potential TMHs and signal peptides (SPs) in MscS2, we used TOPCONS and DeepTMHMM. However, the results were inconclusive, as the tools suggest the presence of one or more TMHs for only 13 MscS2 proteins while predicting no TMH for eight MscS 2 and contradicting each other on the remaining sequences (**Fig 1A**). Furthermore, the tools predict a SP for 25-50% of the sequences. However, TMHs are often mistaken for SPs in these kinds of analyses (49,50). Additionally, for some MscS2 the sequence length just before the MscS β-domain seems insufficient to accommodate two TMHs which are typically at least 17 aa long (51,52). For example, the sequence length of Enterocytozoonidae and close neighbors, as well as *Trachipleistophora hominis* MscS2 range from 73-95 aa (**Fig 1B**) leaving only 22-45 aa for one or two potential TMHs and a connecting loop. Taken together, based on primary sequence it remains elusive how many TMHs microsporidian MscS2 have.

**Fig 1.**
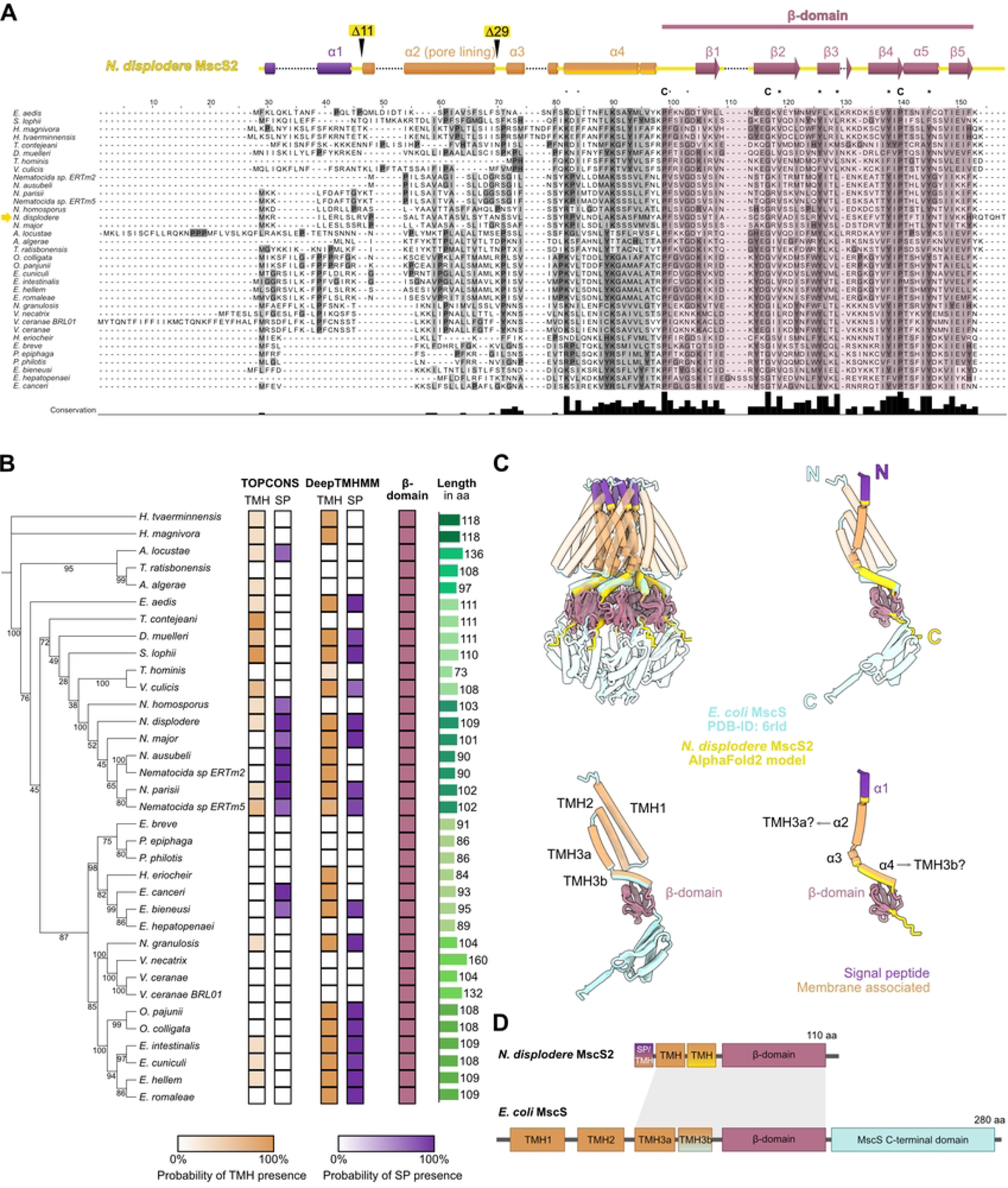
MscS2 is highly conserved among microsporidia and drastically shortened to the minimally required domains. **A)** Sequence alignment of MscS2 from all sequenced microsporidian species and indicated secondary structure elements of *Nematocida displodere* MscS2 with truncation marks. The conserved MscS β-domain is highlighted in shades of pinkish red. Residues marked with “C” are fully conserved, residues with “*” are conserved between groups with very similar characteristics and residues with “.” are conserved between groups with weakly similar characteristics. Multiple sequence alignment was done using ClustalW (38) with default settings and colored according to conservation with “Clustal” in Jalview (v2.11.2.6) and set to black and white. α, alpha-helix; β, beta-strand; MscS, mechanosensitive ion channel. **B)** Cladogram of microsporidia MscS2 with indication of the presence of one or more transmembrane helices (TMH), a signal peptide (SP), the MscS β-domain, and the amino acid sequence length. **C)** Structural comparison and superimposition *N. displodere* MscS2 Alphafold2 model with *E. coli* MscS (PDB-ID 6RLD) channel and single chain. N, N-terminus; C, Carboxy-terminus; TMH, transmembrane helix. All membrane interacting structures are indicated in orange, all signal peptides are colored violet and all MscS β-domains are marked in pinkish red. **D)** Schematic domain architecture of *N. displodere* MscS2, representative for microsporidia, and E. coli MscS, representing bacteria. The grey area indicates the conserved region.

Moving from primary sequence to 3D protein structure, we created an AlphaFold model of *N. displodere* MscS2. While the model is of high confidence in the MscS β-domain and the adjacent helix, the protein fold and domain orientation of the remaining N-terminus however are of low confidence (**S2B Fig**). A hydropathicity prediction of the N-terminal MscS2 sequence (**S2C Fig**) indicates the presence of hydrophobic residues suggesting a transmembrane region. To assess structural similarities and differences of MscS2 to a bacterial homolog, we compared the AlphaFold model to the structure of *Ec*MscS. A superimposition of both proteins highlights the conserved MscS β-domain fold and indicates that the MscS2 helices might correspond to TMH3a (pore-lining) and TMH3b in the bacterial homolog (**Fig 1C**). In *Ec*MscS TMH3 was stated to be essential for channel gating function (32,53). To summarize, microsporidian MscS2 is heavily compacted and lacks N-terminal TMH1 and possibly TMH2 which, based on shorter size, might correspond to a signal sequence. However, with TMH3 and the conserved β-domain, the microsporidian MscS2 maintains the essential domains of a functional mechanosensitive ion channel (**Fig 1D**).

### *Nematocida displodere* MscS2 forms a 400-kDa assembly

Next, we used the obtained information on structure prediction and architecture to design constructs for recombinant MscS2 production. To produce MscS2 and analyze it via single-particle cryo-EM, we ordered codon-optimized *mscS2* from four different microsporidian species: *Nematocida displodere*, *Encephalitozoon cuniculi*, *A. locustae* and *Enterocytozoon bieneusi.* Out of those, *mscS2* from *N. displodere* produced the highest amount of protein (**S3A Fig**) and was used for large-scale protein production and purification. Based on our bioinformatics analyses, *N. displodere* MscS2 (hereafter referred to as “MscS2” or “*mscS2*”) is a transmembrane protein, with one or two transmembrane helices, depending on whether the first is a signal peptide (SP) or not. We thus designed three different *mscS2* constructs, codon-optimized for production in *E. coli* cells: A full-length version of *mscS2*, and two N-terminally truncated versions, *mscS2Δ2-11* and *mscS2Δ2-29* which lack part of, or the whole predicted SP sequence, respectively.

We performed initial solubilization trials of MscS2 variants indicating DDM and LMNG to be the most effective detergents for solubilization. The purifications of pure MscS2 and MscS2Δ2-11 were not successful due degradation and significant aggregation after solubilization of both variants. In addition, the amounts of these two variants were substantially lower than that of MscS2Δ2-29, which could readily be purified to homogeneity in both DDM and LMNG. Initial size-exclusion chromatography (SEC) analysis of the purified His14-SUMO-MscS2Δ2-29 shows a main peak corresponding to a predicted molecular mass of around 320 kDa (according to elution volume), both when solubilized with DDM and LMNG (**S4A Fig**). After tag cleavage, the SEC profile shows a single symmetric peak which corresponds to an even higher molecular weight of circa 430 kDa (**Fig 2A and S4A Fig**). This indicates that the His14-SUMO tag poses a steric hindrance at the N-terminus, thereby preventing higher oligomer formation. The predicted mass from SEC is > 6-fold higher than the theoretical weight of 64 kDa for the expected homo-heptamer (lacking the weight from the detergent micelle). Visualization of the SEC peak fractions via SDS-PAGE indicates a single protein band corresponding to MscS2Δ2-29 monomers (**Fig 2A and S4B Fig**), suggesting that MscS2Δ2-29 oligomerize and form a defined high molecular weight protein-detergent complex. To assess whether this assembly is concentration-dependent, we diluted the protein sample and analyzed it using mass photometry. The mass photometry analysis showed a main peak corresponding to 400 kDa (28 nM) and 375 kDa (56 nM) (**Fig 2B**). The peaks of low molecular weight correspond to background noise **(S4F Fig)**. This indicates that the assembly remains stable even when diluted and confirms the unexpectedly high mass indicated by the SEC.

**Fig 2.**
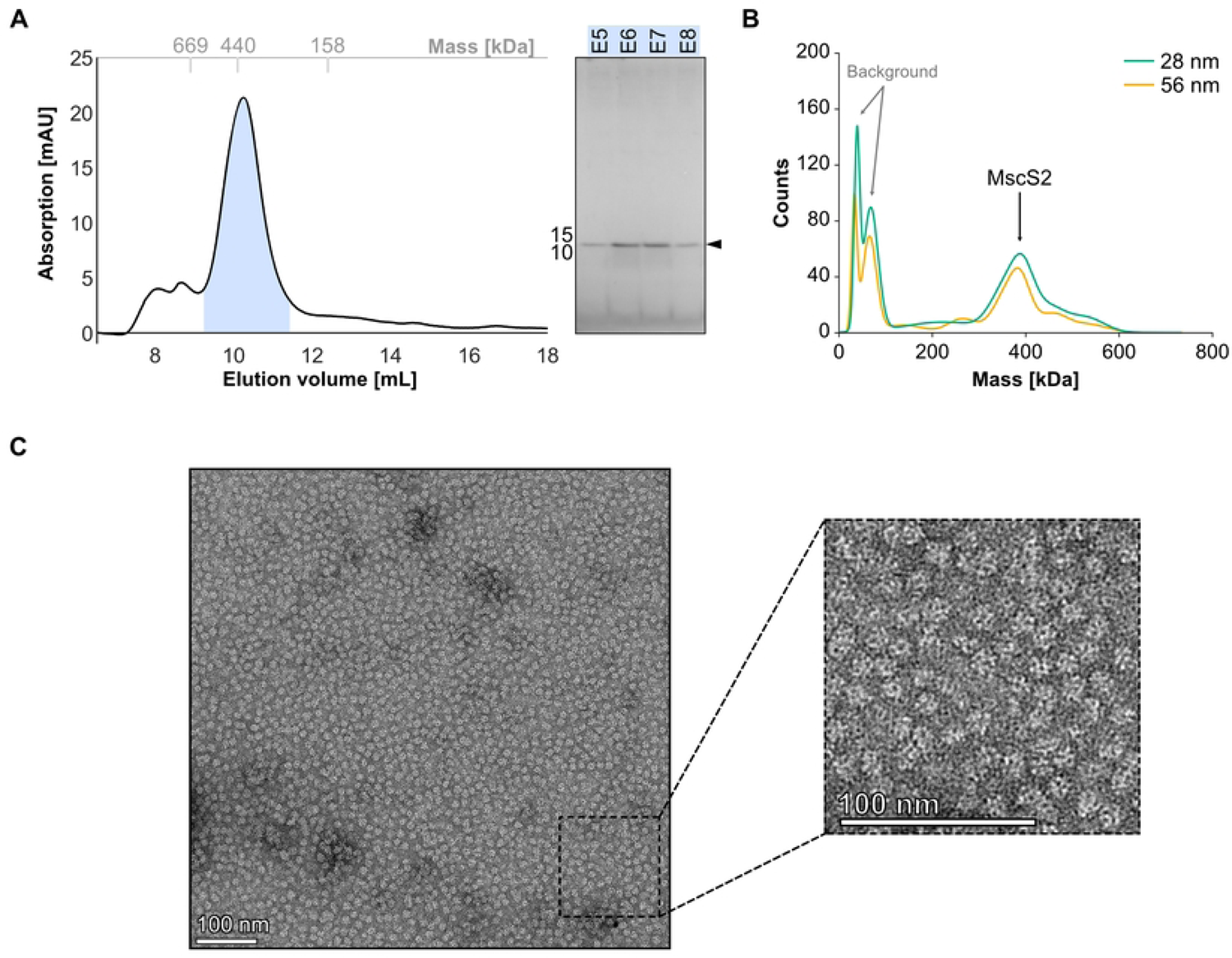
*Nematocida displodere* MscS2Δ2-29 assemble into a high-molecular weight complex. **A)** Size exclusion chromatography (SEC) elution profile of MscS2Δ2-29 [15µM] sample on a Superdex 200 increase 10/300 GL column with SDS-PAGE analysis of the main peak fractions highlighted by a blue shade. The x-axis of the SEC profile shows the elution volume in milliliters (mL), the y-axis displays the absorption in milli-absorbance units and the top x-axis (grey) indicates the molecular weight of standard globular proteins. **B)** Mass photometry analysis of MscS2Δ2-29 indicating a molecular weight of 400 (±60) kDa and 375 (±30) kDa for MscS2Δ2-29 at concentrations 28 nM and 56 nM, respectively. **C)** Representative transmission electron microscopy (TEM) micrograph of negative-stained MscS2Δ2-29 particles at with zoom into the lower right corner. Scale bar diameter is 100 nm.

To gain more insight into the large oligomeric assembly we performed negative-stain transmission electron microscopy (TEM) to assess particle abundance and purity. Confirming the high mass observed in SEC and mass photometry, the particles were significantly larger and different in shape (**Fig 2C**) compared to MscS particles from *E. coli* visualized via TEM (56), suggesting an unusual oligomeric state.

### MscS2 forms a homo-heptameric channel that assembles into a six-channel super structure

While the negative stain micrographs do not show the expected heptameric pore-shaped particles, they confirm the presence of a homogeneous, large protein complex as seen in the SEC and mass photometry analysis. To further characterize this high molecular weight complex formed by MscS2Δ2-29, we collected a single-particle data set using cryo-EM. We observed similarly shaped particles (**Fig 3A**) as before with negative stain EM (**Fig 2C**). A 2D classification of 300,757 particles resulted in unusual, wind-mill-shaped 2D class averages (**Fig 3B**). Further consensus 3D refinement without symmetry led to an 8.3 Å volume. The determined volume resembles an asymmetric, flexible six-way cross-joint (**Fig 3C**). There, six individual channels are oriented in a heterogeneous manner to one another with the N-termini facing inwards (**Fig 3D**) and oligomerizes via their transmembrane domains. We were unable to achieve a higher resolution due to inherent flexibility within the protein complex, with the different heptamers having different tilts and rotations (**Fig 3C**). The tight packing suggests that the presence of the first 29 amino acids, which were truncated to remove the predicted SP and to obtain a homogeneous protein sample, would not allow MscS2 to form this unusual assembly. However, we assume that the only stable condition of the truncated MscS2 channels *in vitro* is this 6-channel superstructure, as electron microscopy and mass-determination experiments are dominated by the large oligomeric complex.

**Fig 3.**
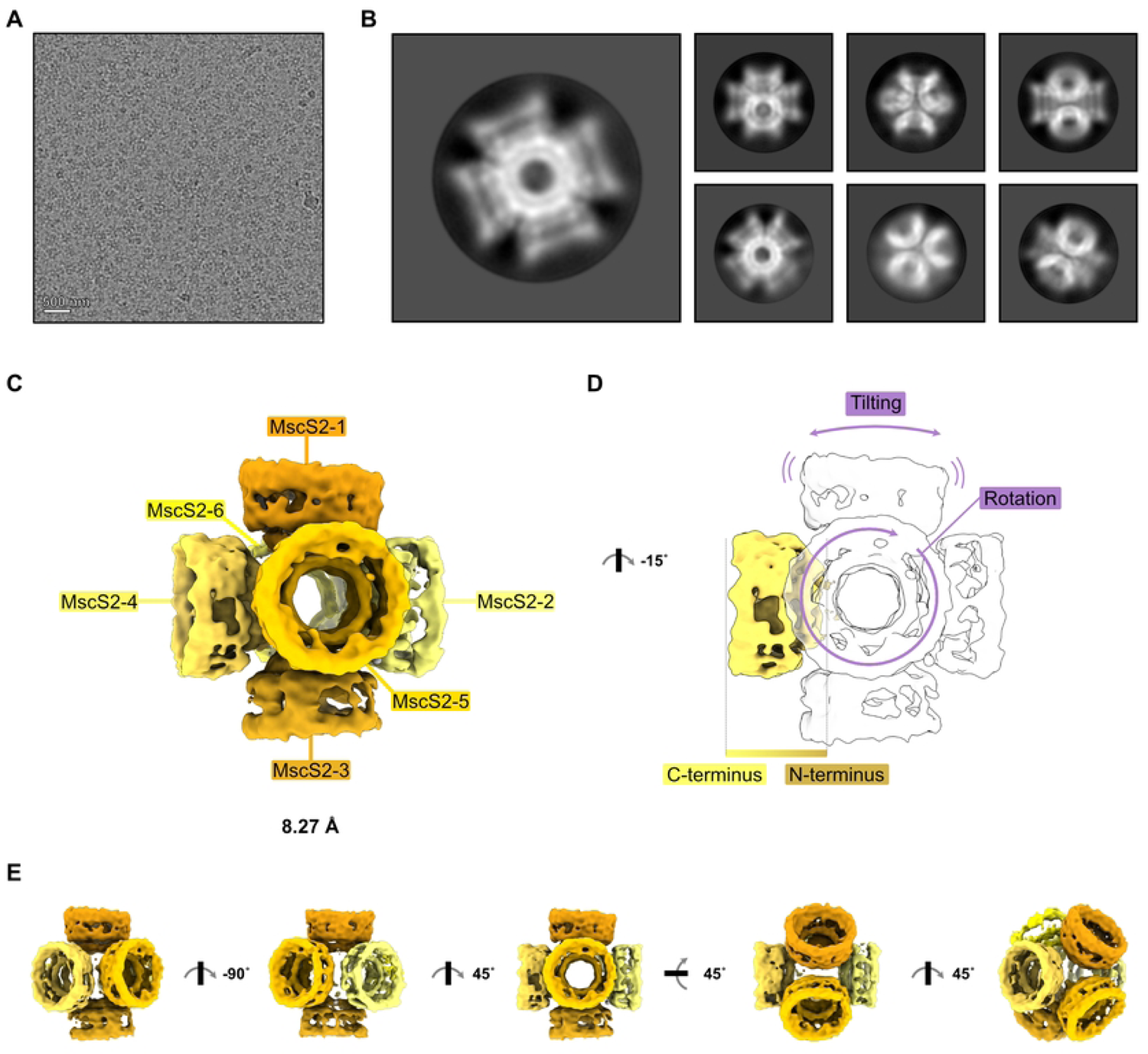
*Nematocida displodere* MscS2Δ2-29 assembles into a six-channel superstructure *in vitro*. **A)** Representative cryo-EM micrograph of the MscS2Δ2-29 sample. **B)** Representative 2D class averages of MscS2Δ2-29 particles. **C)** MscS2Δ2-29 cryo-EM density colored in shades of yellow with labeled heptamer channels and resolution in Å. **D)** Rotationally related view of panel C) with one channel colored to visualize the directionality, N-terminus oriented towards the center and the C-terminus outwards. Heterogeneity and flexibility of individual MscS2Δ2-29 channels are indicated with purple arrows. **E)** Rotationally related views of the MscS2Δ2-29 super structure.

Next, we extracted one subunit of the 6-channel superstructure and superimposed it with the predicted AlphaFold model of the heptameric, full-length MscS2 (with the first 29-aa hidden) (**Fig 4A**) to assess its individual oligomeric state. The AlphaFold homo-heptamer fits into the obtained cryo-EM density, supporting the hypothesis that *N. displodere* MscS2 indeed forms a homo-heptamer, like the bacterial homolog (54). The MscS2 volume has a pore size of ∼30 Å, measured within the MscS β-domain, an outer diameter of ca. 65 Å and is about 40 Å in length along its central axis. To compare it to the bacterial homolog, we superimpose it with an 8 Å low-pass filtered map of *Ec*MscS (PDB id: 6RLD) (**Fig 4B left panel**). The compaction of MscS2 becomes very evident as it only fills out one-third of the filtered *Ec*MscS density, even considering that one potential TMH was removed. However, the MscS2 volume overlays nicely with the “middle section” of *Ec*MscS, the MscS β-domain as well as the amphipathic TMH3b (**Fig 4B right panel**). In *Ec*MscS, TMH3b has been shown to interact with the cytoplasmic domain and is essential for channel gating function (32). It is therefore reasonable to assume that MscS2 has the prerequisites for a functional gating mechanism. How MscS2 embeds in the membrane bilayer to sense membrane tension, however, could not be determined in this study. While in *Ec*MscS, TMH1 and TMH2 pass through the membrane bilayer in an antiparallel manner and act as tension sensor (55,56), the *N. displodere* MscS2 N-terminus is not sufficient to pass through the membrane twice, even when including the missing 29 amino acids. Thus, it is likely that the membrane interaction, involved lipids, and possibly the sensing mechanism, differ between *Ec*MscS and microsporidian MscS2. Another difference between the two channels is the missing C-terminus in MscS2 (2) which is part of the cytoplasmic domain in *Ec*MscS (57). It harbors the α/β-domain and the β-barrel which confer ion selectivity in MscS (57,58). *Ec*MscS variants lacking the β-barrel, remained active under hypoosmotic shock but were less abundant or less stable in the membranes compared to wild-type MscS, while *Ec*MscS variants missing both domains were not even inserted into *E. coli* membranes (33). It is therefore puzzling how microsporidia would employ such a reduced version of MscS2. One possibility is that MscS2 in microsporidia do not have any ion selectivity. Further, depending on where the channel is located within the microsporidian cell, different lipid interactions, as compared to *Ec*MscS, could account for a different sensing mechanism.

**Fig 4.**
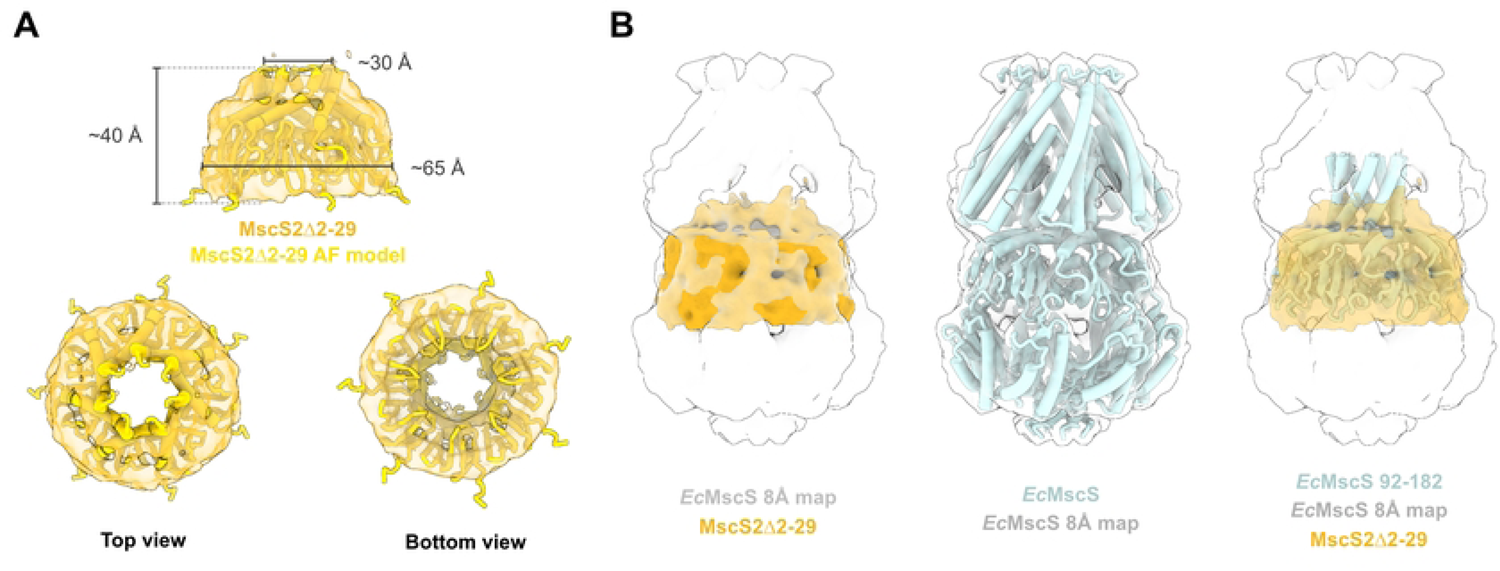
*Nematocida displodere* MscS2 cryo-EM volume embeds well in the MscS β-region of *Ec*MscS. **A)** Top panel: Side view of *N. displodere* MscS2Δ2-29 electron density (goldenrod transparent) superimposed with the corresponding AlphaFold model (gold). Approximate inner and outer diameter, and height of the electron density map (σ-level 0.006) are indicated. Bottom panel: Top and bottom view of the cryo-EM density superimposed with the AlphaFold model. **B)** *E. coli* MscS (6RLD) low-pass filtered electron density map (lightgray transparent) superimposed with the MscS2Δ2-29 volume (left panel), *Ec*MscS structure in the 8 Å map (middle) and a truncated version of *Ec*MscS superimposed with the MscS2Δ2-29 channel volume within the *Ec*MscS 8 Å map.

*mscS2* are, as previously mentioned, present in all microsporidia and transcriptional data shows that it is expressed and therefore likely functional (**S5 Fig**). Transcriptome analyses of *Edhazardia aedis*, *Vavraia culicis* (59), *Encephalitozoon cuniculi* (60) and *Nematocida parisii* (1) suggest that *mscS2* is predominantly upregulated during proliferative stages, such as the schizogony and merogony, but also moderately expressed during stages that require morphological changes, like sporogony, sporulation and nuclear dissociation (**Fig S5**). We therefore assume that MscS2 might mimic the function of the bacterial MscS and regulate homeostasis. However, further experiments are necessary to elucidate the detailed structure and function of the protein. Suggestions for future research include raising antibodies against MscS2, which could aid in locating the channel in infected *C. elegans* as well as pinpointing at which stage of the microsporidian lifecycle MscS2 is produced. Additionally, such antibodies could be used to stabilize the channel, and aid in both purification and especially structure determination. Further, it will be exciting to uncover how microsporidia can operate this minimal version of MscS2, regarding the sensing and gating mechanism.

To conclude, our findings show that *N. displodere* MscS2 forms a homo-heptameric channel that retains the MscS β-domain and TMH3b, which are paramount for channel function. We also show a remarkable complex assembly, which despite not being physiologically relevant, provides fascinating information on the MscS2 assembly as well as of how membrane proteins can oligomerize when not restricted to the lipid membrane.

## Acknowledgments

The authors thank Prof. Robert Hirt and Dr. Josy ter Beek for interesting discussions regarding the project. We thank Dr. Michael Hall and Camilla Holmlund for help with cryo-EM data collection and Dennis Svedberg for aid with the cladogram. The electron microscopy data was collected at the Umeå Core Facility for electron Microscopy, a node of the Cryo-EM Swedish National Facility, funded by the Knut and Alice Wallenberg, Family Erling Persson and Kempe Foundations, SciLifeLab, Stockholm University, and Umeå University.

## Funding

This work was supported by grants from the Swedish Research Council (2016-03599 & 2023-02423 to R.P-A.B and 2019-02011 to J.B.), Knut and Alice Wallenberg Foundation (R.P-A.B), the European Research Council (ERC Starting Grant PolTube 948655 to J.B.), the SciLifeLab National Fellows program, and MIMS. Open access funding provided by Umea University.

## Data availability

The cryo-EM density volume has been deposited in the EM Data Bank with accession code EMD-19937.

## Supporting Information

**S1 Fig.** Cryo-EM data collection and processing. Initially, 2000 micrographs were collected and manually inspected to remove any affected by drift, poor CTF fits, or low-quality ice, resulting in a total of 1856 micrographs. Particles were picked through Laplacian auto picking and underwent one round of 2D classification to eliminate picking contaminants. From the resulting 300,757 particles, an initial model was generated, and a subsequent consensus refinement, without symmetry, followed by post processing yielded an 8.3 Å volume.

**S2 Fig.** Multiple sequence alignment of *E. coli* MscS and 35 microsporidian MscS2, *N. displodere* MscS2 AlphaFold2-model and hydropathicity prediction. A) Multiple sequence alignment (MSA) of *E. coli* MscS and microsporidian MscS2 shows high conservation of the MscS β-domain and moderate conservation of the transmembrane helix 3b (TMH3b). MSA was generated using ClustalW (38) with default settings, colored according to conservation with “Clustal” in Jalview (v2.11.2.6) and set to black and white. **B)** AlphaFold2 (v2.3.1) prediction of homoheptameric *N. displodere* MscS2 (left), colored by pLDDT confidence measure, with indicated color key, and predicted aligned error plots (right). **C)** Hydropathicity prediction for *N. displodere* MscS2 to identify hydrophobic regions, calculated using ProtScale on the ExPASy server (45).

**S3 Fig.** Test production of full-length MscS2 and purification of truncated MscS2. A) Western Blot analysis of induced and non-induced cultures comparing MscS2 yields from *A. locustae*, *E. bieneusi*, *E. cuniculi* and *N. displodere* produced with a C-terminal His10 tag in *E. coli* Rosetta (DE3). Anti-His6x antibody from mouse was used for immunoprecipitation (1:1000). GFP-MS2-His10 construct served as positive control for protein production and antibody binding. As seen in the blots, the promotor is leaky as MscS2 is produced to some extent in the absence of IPTG. **B)** IMAC purification of DDM-solubilized His14-SUMO-MscS2Δ2-29 visualized by SDS-PAGE showing that the protein can be produced in high amounts. **C)** SDS-PAGE analysis of the fractions corresponding to the reverse IMAC of DDM-solubilized MscS2Δ2-29 post tag cleavage. Fraction used for negative-stain TEM studies is boxed in black and fractions analyzed via SEC are marked with blue boxes. kDa, kilo Dalton; M, marker; IPTG, Isopropyl β-d-1-thiogalactopyranoside; GFP, green-fluorescent protein; MS2, MS2 bacteriophage coat protein; pooled E, pooled elution; FT, flowthrough; SEC, size exclusion chromatography, kDa, kilo Dalton; M, marker; FT, flowthrough; W, wash; E, elution; SUMO, small ubiquitin-related modifier; SEC, size-exclusion chromatography.

**S4 Fig.** Tag cleavage of truncated MscS2 induces an increase in molecular weight indicating higher oligomeric complex formation. A) Size exclusion chromatogram (SEC) profile of DDM and LMNG-solubilized MscS2Δ2-29 with and without (only for DDM) His14-SUMO tag. DDM-solubilized MscS2Δ2-29 elutes at 10.16 mL, His14-SUMO-MscS2Δ2-29 in DDM elutes at 10.87 and the latter solubilized with LMNG elutes at 10.93 mL. The observed main peak shift post tag cleavage is indicated. The x-axis of the SEC profile shows the elution volume in milliliters (mL), the y-axis displays the absorption in milli-absorbance units and a second x-axis at the top shows the molecular weight of standard globular proteins. B-D) SDS-PAGE analyses of the indicated SEC-elution fractions for B) MscS2Δ2-29 (ca. 9 kDa) and C) His14-SUMO-MscS2Δ2-29 (ca. 23 kDa) both solubilized using DDM and D) His14-SUMO-MscS2Δ2-29 solubilized with LMNG. The main SEC peaks in A) correspond to MscS2Δ2-29 with or without His14-SUMO tag. E) Mass photometry analysis comparing the molecular weight of MscS2Δ2-29 with and without His14-SUMO tag. For His14-SUMO-MscS2Δ2-29 (12nM) a main peak corresponding to 158 (± 15.6) kDa was detected and MscS2Δ2-29 (140 nM) produced a main peak at 452 (± 112) kDa. F) Buffer control of the mass photometry experiments showing a peak around 30 (± 4.8) kDa and 61 (± 15.6) kDa.

**S5 Fig.** *mscS2* transcription in different life-cycle stages of *Vavraia culicis*, *Edhazardia aedis*, *Nematocida parisii* and *Encephalitozoon cuniculi*. A,B) Gene expression pattern of *mscS2* in two microsporidians, *Vavraia culicis* (A, VCUG_00285) with a simple, and *Edhazardia aedis* (A, EDEG_01910) with a complex life cycle (59). The life-cycle stages corresponding to the sampling points are indicated. Each developmental stage was sampled in duplicates (59) and is shown in shades of yellow (A) and cyan (B). **C)** Transcript levels of *Nematocida parisii mscS2* (NEPG_00116) over the course of a full infection cycle: 1) 8 hours post infection (hpi) sporoplasm stage, 2) 16 hpi early meront stage, 3) 30 hpi late meront stage, 4) 40 hpi onset of spore formation and 5) 64 hpi spores within membrane-bound vesicles. Samples from 8, 16 and 30 hpi are reported to be dominated by proliferating meronts, and later samples at 40 and 64 hpi harbor a mixture of meront, sporont and mature spore stages. Developmental stages were assessed by differential interference contrast microscopy and fluorescence in situ hybridization (1). **D)** Transcriptional profile of mscS2 in *Encephalitozoon cuniculi* (Ecu09_0470) after 24, 48 and 72 hpi. Transcriptomic data indicates that at 24 hpi proliferation rates are throttled due to downregulation of housekeeping genes. Further, at 48 hpi meronts start producing spore-related genes but spore formation is not expected until after 72 hpi (60).

**S1 Table.** Primers used in this study.

**S2 Table.** Cryo-EM data collection and processing.

## Notes

### Competing Interest Statement

The authors have declared no competing interest.

## References

1. Cuomo CA, Desjardins CA, Bakowski MA, Goldberg J, Ma AT, Becnel JJ, et al. Microsporidian genome analysis reveals evolutionary strategies for obligate intracellular growth. Genome Res. 2012 Dec;22(12):2478–88.

2. Nakjang S, Williams TA, Heinz E, Watson AK, Foster PG, Sendra KM, et al. Reduction and Expansion in Microsporidian Genome Evolution: New Insights from Comparative Genomics. Genome Biology and Evolution. 2013 Dec 1;5(12):2285–303.

3. Capella-Gutiérrez S, Marcet-Houben M, Gabaldón T. Phylogenomics supports microsporidia as the earliest diverging clade of sequenced fungi. BMC Biol. 2012 Dec;10(1):47.

4. Keeling PJ, Slamovits CH. Simplicity and Complexity of Microsporidian Genomes. Eukaryotic Cell. 2004 Dec;3(6):1363.

5. Keeling PJ, Fast NM. Microsporidia: Biology and Evolution of Highly Reduced Intracellular Parasites. Annual Review of Microbiology. 2002;56(1):93–116.

6. Didier ES. Microsporidiosis: an emerging and opportunistic infection in humans and animals. Acta Trop. 2005 Apr;94(1):61–76.

7. Han B, Takvorian PM, Weiss LM. Invasion of Host Cells by Microsporidia. Front Microbiol. 2020 Feb 18;11:172.

8. Stentiford GD, Becnel J.J., Weiss LM, Keeling PJ, Didier ES, Williams B. AP, et al. Microsporidia – Emergent Pathogens in the Global Food Chain. Trends in Parasitology. 2016 Apr;32(4):336–48.

9. Huang WF, Solter LF, Yau PM, Imai BS. Nosema ceranae Escapes Fumagillin Control in Honey Bees. Schneider DS, editor. PLoS Pathog. 2013 Mar 7;9(3):e1003185.

10. Li JH, Evans JD, Li WF, Zhao YZ, DeGrandi-Hoffman G, Huang SK, et al. New evidence showing that the destruction of gut bacteria by antibiotic treatment could increase the honey bee’s vulnerability to Nosema infection. Rueppell O, editor. PLoS ONE. 2017 Nov 10;12(11):e0187505.

11. Didier ES, Weiss LM. Microsporidiosis: current status. Current Opinion in Infectious Diseases. 2006 Oct;19(5):485–92.

12. Didier ES, Weiss LM. Microsporidiosis: not just in AIDS patients. Current Opinion in Infectious Diseases. 2011 Oct;24(5):490.

13. Huang Q, Chen J, Lv Q, Long M, Pan G, Zhou Z. Germination of Microsporidian Spores: The Known and Unknown. JoF. 2023 Jul 22;9(7):774.

14. Pleshinger J, Weidner E. The microsporidian spore invasion tube. IV. Discharge activation begins with pH-triggered Ca2+ influx. The Journal of cell biology. 1985 Jun 1;100(6):1834–8.

15. Weidner E. Ultrastructural study of microsporidian invasion into cells. Zeitschrift für Parasitenkunde. 1972 Sep 1;40(3):227–42.

16. Chang R, Davydov A, Jaroenlak P, Budaitis B, Ekiert DC, Bhabha G, et al. Energetics of the Microsporidian Polar Tube Invasion Machinery [Internet]. Biophysics; 2023 Jan [cited 2024 Mar 7]. Available from: http://biorxiv.org/lookup/doi/10.1101/2023.01.17.524456

17. Dean P, Sendra KM, Williams TA, Watson AK, Major P, Nakjang S, et al. Transporter gene acquisition and innovation in the evolution of Microsporidia intracellular parasites. Nat Commun. 2018 Apr 27;9(1):1709.

18. Han B, Ma Y, Tu V, Tomita T, Mayoral J, Williams T, et al. Microsporidia Interact with Host Cell Mitochondria via Voltage-Dependent Anion Channels Using Sporoplasm Surface Protein 1. Sibley LD, editor. mBio. 2019 Aug 27;10(4):e01944–19.

19. Dean P, Hirt RP, Embley TM. Microsporidia: Why Make Nucleotides if You Can Steal Them? Gubbels MJ, editor. PLoS Pathog. 2016 Nov 17;12(11):e1005870.

20. Antao NV, Lam C, Davydov A, Riggi M, Sall J, Petzold C, et al. 3D reconstructions of parasite development and the intracellular niche of the microsporidian pathogen Encephalitozoon intestinalis. Nat Commun. 2023 Nov 23;14(1):7662.

21. Vivarès C. Functional and evolutionary analysis of a eukaryotic parasitic genome. Current Opinion in Microbiology. 2002 Oct 1;5(5):499–505.

22. Tamim El Jarkass H, Reinke AW. The ins and outs of host-microsporidia interactions during invasion, proliferation and exit. Cellular Microbiology [Internet]. 2020 Nov [cited 2024 Mar 7];22(11). Available from: https://onlinelibrary.wiley.com/doi/10.1111/cmi.13247

23. Timofeev S, Tokarev Y, Dolgikh V. Energy metabolism and its evolution in Microsporidia and allied taxa. Parasitol Res. 2020 May;119(5):1433–41.

24. Jespersen N, Monrroy L, Barandun J. Impact of Genome Reduction in Microsporidia. In: Weiss LM, Reinke AW, editors. Microsporidia [Internet]. Cham: Springer International Publishing; 2022 [cited 2023 Sep 10]. p. 1–42. (Experientia Supplementum; vol. 114). Available from: https://link.springer.com/10.1007/978-3-030-93306-7_1

25. Pombert JF, Haag KL, Beidas S, Ebert D, Keeling PJ. The Ordospora colligata Genome: Evolution of Extreme Reduction in Microsporidia and Host-To-Parasite Horizontal Gene Transfer. Boothroyd JC, editor. mBio. 2015 Feb 27;6(1):e02400–14.

26. Corradi N. Microsporidia: Eukaryotic Intracellular Parasites Shaped by Gene Loss and Horizontal Gene Transfers. Annu Rev Microbiol. 2015 Oct 15;69(1):167–83.

27. Kung C, Martinac B, Sukharev S. Mechanosensitive Channels in Microbes. Annu Rev Microbiol. 2010 Oct 13;64(1):313–29.

28. Zhang Y, Daday C, Gu RX, Cox CD, Martinac B, De Groot BL, et al. Visualization of the mechanosensitive ion channel MscS under membrane tension. Nature. 2021 Feb 18;590(7846):509–14.

29. Nakayama Y, Yoshimura K, Iida H. Organellar mechanosensitive channels in fission yeast regulate the hypo-osmotic shock response. Nat Commun. 2012 Aug 21;3(1):1020.

30. Hurst AC, Petrov E, Kloda A, Nguyen T, Hool L, Martinac B. MscS, the bacterial mechanosensitive channel of small conductance. The International Journal of Biochemistry & Cell Biology. 2008;40(4):581–5.

31. Edwards MD, Li Y, Kim S, Miller S, Bartlett W, Black S, et al. Pivotal role of the glycine-rich TM3 helix in gating the MscS mechanosensitive channel. Nat Struct Mol Biol. 2005 Feb;12(2):113–9.

32. Wang X, Tang S, Wen X, Hong L, Hong F, Li Y. Transmembrane TM3b of Mechanosensitive Channel MscS Interacts With Cytoplasmic Domain Cyto-Helix. Front Physiol. 2018 Oct 1;9:1389.

33. Schumann U, Edwards MD, Li C, Booth IR. The conserved carboxy-terminus of the MscS mechanosensitive channel is not essential but increases stability and activity. FEBS Letters. 2004 Aug 13;572(1–3):233–7.

34. Svedberg D, Winiger RR, Berg A, Sharma H, Tellgren-Roth C, Debrunner-Vossbrinck BA, et al. Functional annotation of a divergent genome using sequence and structure-based similarity. BMC Genomics. 2024 Jan 2;25(1):6.

35. Corradi N, Akiyoshi DE, Morrison HG, Feng X, Weiss LM, Tzipori S, et al. Patterns of Genome Evolution among the Microsporidian Parasites Encephalitozoon cuniculi, Antonospora locustae and Enterocytozoon bieneusi. Butler G, editor. PLoS ONE. 2007 Dec 5;2(12):e1277.

36. Slamovits CH, Fast NM, Law JS, Keeling PJ. Genome Compaction and Stability in Microsporidian Intracellular Parasites. Current Biology. 2004 May;14(10):891–6.

37. Nordberg H, Cantor M, Dusheyko S, Hua S, Poliakov A, Shabalov I, et al. The genome portal of the Department of Energy Joint Genome Institute: 2014 updates. Nucl Acids Res. 2014 Jan;42(D1):D26–31.

38. Madeira F, Pearce M, Tivey ARN, Basutkar P, Lee J, Edbali O, et al. Search and sequence analysis tools services from EMBL-EBI in 2022. Nucleic Acids Research. 2022 Jul 5;50(W1):W276–9.

39. Waterhouse AM, Procter JB, Martin DMA, Clamp M, Barton GJ. Jalview Version 2—a multiple sequence alignment editor and analysis workbench. Bioinformatics. 2009 May 1;25(9):1189–91.

40. Capella-Gutiérrez S, Silla-Martínez JM, Gabaldón T. trimAl: a tool for automated alignment trimming in large-scale phylogenetic analyses. Bioinformatics. 2009 Aug 1;25(15):1972–3.

41. Minh BQ, Schmidt HA, Chernomor O, Schrempf D, Woodhams MD, von Haeseler A, et al. IQ-TREE 2: New Models and Efficient Methods for Phylogenetic Inference in the Genomic Era. Molecular Biology and Evolution. 2020 May 1;37(5):1530–4.

42. Nguyen LT, Schmidt HA, von Haeseler A, Minh BQ. IQ-TREE: A Fast and Effective Stochastic Algorithm for Estimating Maximum-Likelihood Phylogenies. Molecular Biology and Evolution. 2015 Jan 1;32(1):268–74.

43. Hallgren J, Tsirigos KD, Pedersen MD, Almagro Armenteros JJ, Marcatili P, Nielsen H, et al. DeepTMHMM predicts alpha and beta transmembrane proteins using deep neural networks [Internet]. Bioinformatics; 2022 Apr [cited 2024 Mar 20]. Available from: http://biorxiv.org/lookup/doi/10.1101/2022.04.08.487609

44. Tsirigos KD, Peters C, Shu N, Käll L, Elofsson A. The TOPCONS web server for consensus prediction of membrane protein topology and signal peptides. Nucleic Acids Res. 2015 Jul 1;43(W1):W401–7.

45. Gasteiger E, Hoogland C, Gattiker A, Duvaud S, Wilkins MR, Appel RD, et al. Protein Identification and Analysis Tools on the ExPASy Server. In: Walker JM, editor. The Proteomics Protocols Handbook [Internet]. Totowa, NJ: Humana Press; 2005 [cited 2024 Mar 19]. p. 571–607. Available from: http://link.springer.com/10.1385/1-59259-890-0:571

46. Kyte J, Doolittle RF. A simple method for displaying the hydropathic character of a protein. Journal of Molecular Biology. 1982 May;157(1):105–32.

47. Mirdita M, Schütze K, Moriwaki Y, Heo L, Ovchinnikov S, Steinegger M. ColabFold: making protein folding accessible to all. Nat Methods. 2022 Jun;19(6):679–82.

48. Jumper J, Evans R, Pritzel A, Green T, Figurnov M, Ronneberger O, et al. Highly accurate protein structure prediction with AlphaFold. Nature. 2021 Aug;596(7873):583–9.

49. Reynolds SM, Käll L, Riffle ME, Bilmes JA, Noble WS. Transmembrane Topology and Signal Peptide Prediction Using Dynamic Bayesian Networks. Rost B, editor. PLoS Comput Biol. 2008 Nov 7;4(11):e1000213.

50. Lao DM, Arai M, Ikeda M, Shimizu T. The presence of signal peptide significantly affectstransmembrane topology prediction. Bioinformatics. 2002 Dec 1;18(12):1562–6.

51. Hildebrand PW, Preissner R, Frömmel C. Structural features of transmembrane helices. FEBS Letters. 2004 Feb 13;559(1–3):145–51.

52. Heijne G. Membrane Proteins. The Amino Acid Composition of Membrane-Penetrating Segments. Eur J Biochem. 1981 Nov;120(2):275–8.

53. Martinac B. Structural plasticity in MS channels. Nat Struct Mol Biol. 2005 Feb;12(2):104–5.

54. Bass RB, Strop P, Barclay M, Rees DC. Crystal Structure of *Escherichia coli* MscS, a Voltage-Modulated and Mechanosensitive Channel. Science. 2002 Nov 22;298(5598):1582–7.

55. Pliotas C, Dahl ACE, Rasmussen T, Mahendran KR, Smith TK, Marius P, et al. The role of lipids in mechanosensation. Nat Struct Mol Biol. 2015 Dec;22(12):991–8.

56. Deng Z, Maksaev G, Schlegel AM, Zhang J, Rau M, Fitzpatrick JAJ, et al. Structural mechanism for gating of a eukaryotic mechanosensitive channel of small conductance. Nat Commun. 2020 Jul 23;11(1):3690.

57. Koprowski P, Kubalski A. C Termini of the Escherichia coli Mechanosensitive Ion Channel (MscS) Move Apart upon the Channel Opening. Journal of Biological Chemistry. 2003 Mar;278(13):11237–45.

58. Zhang X, Wang J, Feng Y, Ge J, Li W, Sun W, et al. Structure and molecular mechanism of an anion-selective mechanosensitive channel of small conductance. Proc Natl Acad Sci USA. 2012 Oct 30;109(44):18180–5.

59. Desjardins CA, Sanscrainte ND, Goldberg JM, Heiman D, Young S, Zeng Q, et al. Contrasting host–pathogen interactions and genome evolution in two generalist and specialist microsporidian pathogens of mosquitoes. Nat Commun. 2015 May 13;6(1):7121.

60. Grisdale CJ, Bowers LC, Didier ES, Fast NM. Transcriptome analysis of the parasite Encephalitozoon cuniculi: an in-depth examination of pre-mRNA splicing in a reduced eukaryote. BMC Genomics. 2013 Dec;14(1):207.

